# Investigating the impact of combination phage and antibiotic therapy: a modeling study

**DOI:** 10.1101/2020.01.08.899476

**Authors:** Selenne Banuelos, Hayriye Gulbudak, Mary Ann Horn, Qimin Huang, Aadrita Nandi, Hwayeon Ryu, Rebecca Segal

## Abstract

Antimicrobial resistance (AMR) is a serious threat to global health today. The spread of AMR, along with the lack of new drug classes in the antibiotic pipeline, has resulted in a renewed interest in phage therapy, which is the use of bacteriophages to treat pathogenic bacterial infections. This therapy, which was successfully used to treat a variety of infections in the early twentieth century, had been largely dismissed due to the discovery of easy to use antibiotics. However, the continuing emergence of antibiotic resistance has motivated new interest in the use of phage therapy to treat bacterial infections. Though various models have been developed to address the AMR-related issues, there are very few studies that consider the effect of phage-antibiotic combination therapy. Moreover, some of biological details such as the effect of the immune system on phage have been neglected. To address these limitations, we utilized a mathematical model to examine the role of the immune response in concert with phage-antibiotic combination therapy compounded with the effects of the immune system on the phages being used for treatment. We explore the effect of phage-antibiotic combination therapy by adjusting the phage and antibiotics dose or altering the timing. The model results show that it is important to consider the host immune system in the model and that frequency and dose of treatment are important considerations for the effectiveness of treatment. Our study can lead to development of optimal antibiotic use and further reduce the health risks of the human-animal-plant-ecosystem interface caused by AMR.

## 1. Introduction

Antimicrobial resistance (AMR) is a serious threat to global health. The Centers for Disease Control and Prevention (CDC) estimates that at least 2 million people become infected by antibiotic-resistant bacteria and at least 23,000 people die each year as a direct result of these infections, costing the United States $55 billion annually [3]. Infections caused by bacteria are usually treated with antibiotics, however, due to over-prescribing and mis-prescribing, many strains of bacteria have become resistant to currently available antibiotics. A list of antibiotic-resistant pathogens, a catalog of 12 families of bacteria, for which new antibiotics are urgently needed, has been provided by the World Health Organization (WHO) [21]. However, since bacteria evolve resistance to antibiotics at a relatively rapid rate, there has been less commercial interest in developing new antibiotics. Only 6 new antibiotics were approved by the Food and Drug Administration (FDA) for use in the United States from 2010 to 2016, an obvious downward trend compared to the 16 new antibiotics approved by FDA between 1983 and 1987 [41]. In 2015, a global action plan on antimicrobial resistance (GAP-AMR) was endorsed at the World Health Assembly, and one of the five strategic objectives of the GAP-AMR is to optimize the use of antimicrobial agents [65]. In 2018, the U.S. government launched the Antimicrobial Resistance Challenge to call for leaders from around the world to work together to improve antibiotic use, accelerate research on new antibiotics and antibiotic alternatives [3]. The spread of antimicrobial resistance combined with the lack of new drug classes in the antibiotic pipeline has resulted in a resurgence of interest in phage therapy.

Phage therapy is the use of bacteriophages to treat pathogenic bacterial infections. Before the widespread use of antibiotics, phage therapy was successfully applied in treating a variety of infections in the 1920s and 1930s [45]. Due to a poor understanding of the biological nature of phages, medical limitations of the day, and introduction of broader spectrum antibiotics, phage therapy was largely dismissed by most of western medicine in the 1940s [39]. However, the rise of antibiotic resistance has resulted in renewed interest in using phage to treat bacterial infections [52]. One of the first international, single-blind clinical trials of phage therapy, which aimed to target 220 burn patients with wounds infected by *Escherichia coli* or *Pseudomonas aeruginosa*, was launched in 2015 [17, 33, 47]. Furthermore, clinical trials are currently underway to explore phage treatment for infections caused by *Staphylococcus aureus*, particularly for respiratory tract infection (e.g., pneumonia), and to reduce the population of pathogens in ready-to-eat foods and meat [1, 26, 27, 25, 34, 51]. In contrast to antibiotics, bacteria sensitivity to phages is largely specific for both species and strain, which can be considered as a major advantage, since the effects of antibiotics on commensal gut microbes are notorious for secondary outcomes such as antibiotic-associated diarrhea and *C. difficile* infection [46]. See Figure 1 for a timeline of important events in the development and use of phage therapy.

**Fig. 1:**
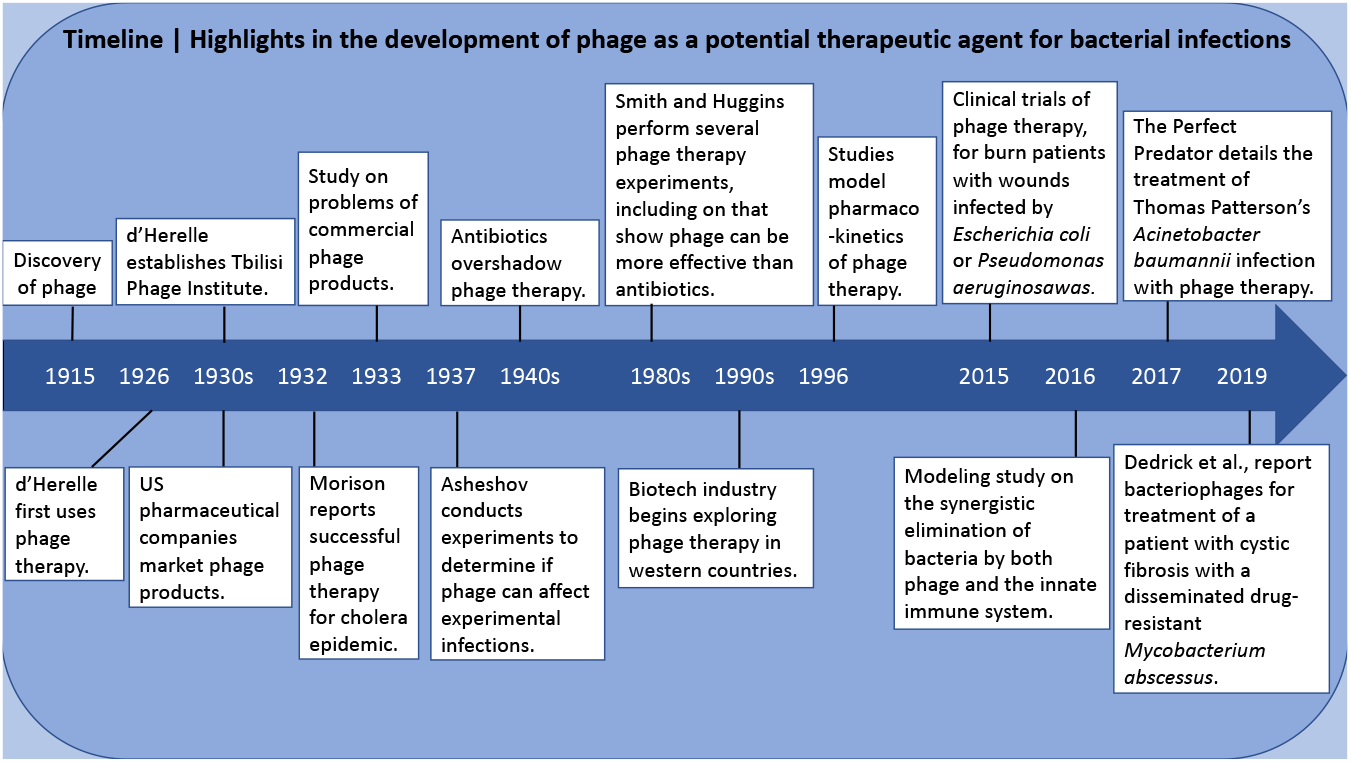
A timeline of important events in the history of phage therapy, adapted and updated from [37].

Because the problem of antibiotic resistant bacteria is complex and growing, with no known solution, various mathematical models have been proposed to explore the dynamics of the variety of systems involved. Most models focus on the transmission dynamics of antibiotic-resistant bacteria at the host population level [4, 5, 7, 8, 9, 10, 11, 14, 18, 19, 24, 28, 31, 32, 30, 40, 58, 59, 60, 61, 62, 63, 20]; some focus on exploring the relative contributions of antibiotics and immune response in the treatment of infection on the bacterial population level [52, 38, 6, 36, 2]. Now, with increasing interest in phage therapy as an alternative or supplement to antibiotic treatment [39], mathematical models incorporating phage therapy have been developed [22, 13, 43, 44, 54, 57, 64, 16, 35, 37, 50, 49]. In particular, Rodriguez-Gonzalez et al. [50] developed a mathematical model of phage-antibiotic combination therapy, representing the interactions among bacteria, phage, antibiotics, and the innate immune system, but ignoring the effect of immune system on phages. Some evidence shows that even though phages are not able to boost innate immunity, bacteria-boosted innate immunity acts against the phages [29], which is an important finding to further explain instances of phage ineffectiveness and to suggest better protocols for using phage therapy. To include this important component, we extended earlier models, in particular, the model developed by Rodriguez-Gonzalez, et al. [50]. The goal is to understand the role of the immune response in concert with phage-antibiotic combination therapy by introducing immune activity related to phages to the model.

We aim to explore the effect of phage-antibiotic combination therapy by adjusting the phage and antibiotic doses and/or altering the timing of the dose(s). Details of the system of nonlinear, ordinary differential equations which take into account the interactions among bacteria, phage, antibiotics, and the immune system are given in Section 2. In Section 3 and 4, equilibria and sensitivity analysis of some reduced cases are pro vided. In Section 5, the simulation results exploring various infection and treatment scenarios are presented, followed by a discussion in Section 6.

## 2. Mathematical Model

We present a deterministic antibiotic-phage combination therapy model that describes density-dependent interactions between two strains of bacteria, phage, antibiotics, and the host immune response. The model development builds on the work by Leung and Weitz [34], and Rodriguez-Gonzalez, et al. [50]. The model in [34] includes phage-sensitive bacteria, phage, and a saturating innate immune response. The phage therapy model in [50] extended [34] to include two bacteria strains, phage therapy, antibiotic treatment and some immune response components. The model presented here adds biological functions not included in these previous models: interactions of the immune response with phage, and the decay of the immune response. See Figure 2 for a schematic diagram of our model.

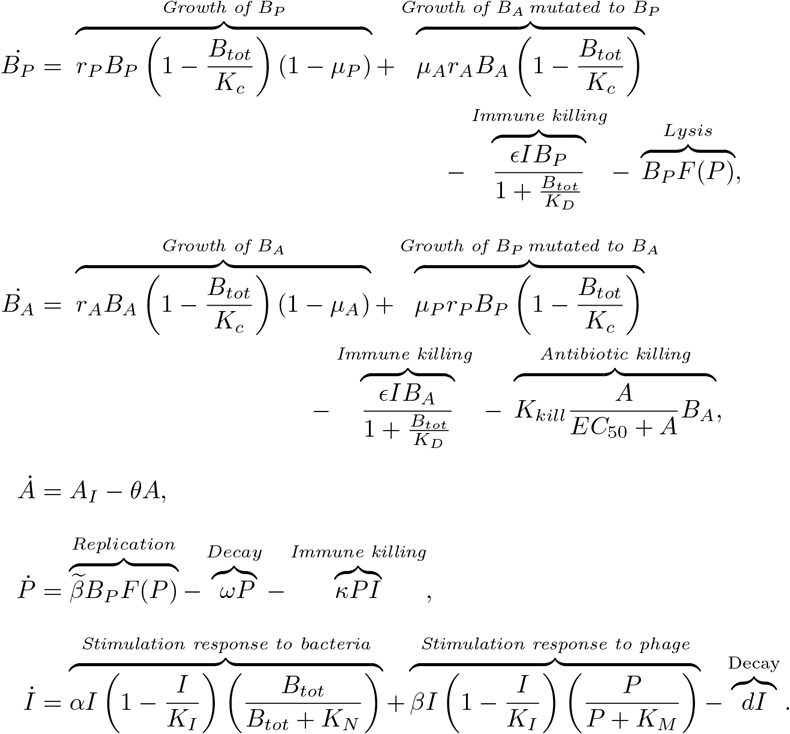

**Fig. 2:**
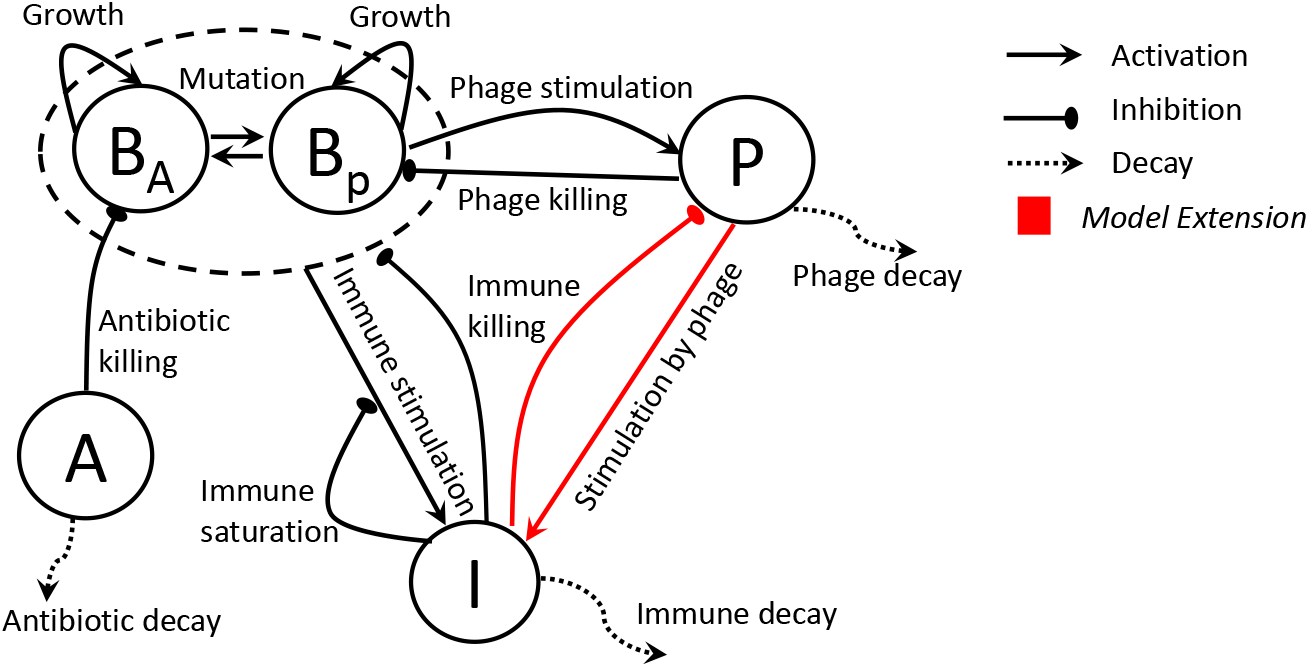
Schematic diagram of the extended phage-antibiotic combination therapy model. Antibiotic-sensitive bacteria (*B*_*A*_) and phage-sensitive bacteria (*B*_*P*_) are targeted by antibiotic (*A*) and the phage (*P*), respectively. The immune response interactions with both bacterial strains are included in the model. In addition, our model extension building on the model in [49] includes the innate immunity (*I*) stimulation by the presence of phage (in red arrows) and the decay of the immune response.

The phage-sensitive (antibiotic-resistant) bacteria, denoted as *B*_*P*_, respond to treatment by phages, *P* whereas the antibiotic-sensitive (phage-resistant) bacteria, *B*_*A*_, responds to treatment by antibiotics, *A*. It can be assumed that bacteria is either sensitive to phages or to antibiotics due to conservation of evolutionary resources in the bacteria [15]. The total immune response (*I*) is activated by the presence of bacteria and phages, and attacks both bacterial strains.

The bacteria grow logistically with growth rate *r*_*i*_, carrying capacity *K*_*c*_, and density dependence *B*_*tot*_ = ∑_*i*_ *B*_*i*_, where *i* ∈ {*A, P*}. We assume that phage-sensitive bacteria mutate to become antibiotic sensitive bacteria with probability *μ*_*P*_. Similarly, *μ*_*A*_ represents the probability of emergence of phage-sensitive mutants from antibiotic-sensitive bacteria. Therefore the growth of the bacteria population is modeled as:

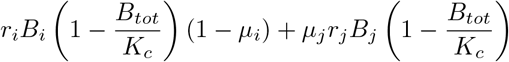

where *i, j* ∈ {A, P} and *i* ≠ *j*. As in [50] both populations of bacteria are killed by an activated innate immune response which includes a density-dependent immune evasion by bacteria. That is, the mass action killing term, ϵ*IB*_*i*_, is scaled by the parameter 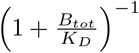. See [34] for more details. The decrease in density of *B*_*A*_ by the antibiotic treatment is approximated by a Hill function as in [50]. The phage-sensitive bacteria are infected and lysed by phage at a rate of *F* (*P*). Following the work in [50] two phage infection modalities, *F* (*P*), are considered - homogeneous mixing and heterogeneous mixing. The homogeneous mixing modality is given by *F* (*P*) = *ϕP* so that the infection rate is proportional to the phage density. The second modality is given by *F* (*P*) = *ϕP*^*γ*^ where *γ* is the power-law exponent. The homogeneous mixing modality is assumed for our analytical results in Section 3 and sensitivity analysis in Section 4, whereas the heterogeneous mixing model is used for the numerical analysis in Section 5. Mathematical analysis is not valid with fractional exponents, but it is likely that the phage distribution is heterogeneous.

We assume that once the antibiotic treatment is administered it is injected at a constant rate where *A*^*^ = *A*_*I*_/*θ*.

The growth in the phage density is due to the release of phage through lytic infection of *B*_*P*_ at a rate of 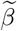. Free phage particles decay at a rate *ω*. One of the novel biological features included in this model is the effect the immune response on the phage virus. The differences in the effectiveness of phage therapy between *in vitro* and *in vivo* suggest that the infected mammalian host’s immune response may be responsible for bacterial phage resistance [29]. The per capita kill rate of phage by the immune response is denoted by *κ*.

As in [34] we assume there is a saturated innate immune response that is activated by bacteria. We have included a saturated innate immune response that is activated by the presence of phage where *β* is the maximum growth rage, *K*_*I*_ is the maximum capacity, and *K*_*M*_ is the phage concentration at which the immune response growth rate is half its maximum. In addition, we assume that *d* is the rate of decay in the immune response. Parameter values are given in Table 1.

**Table 1:**
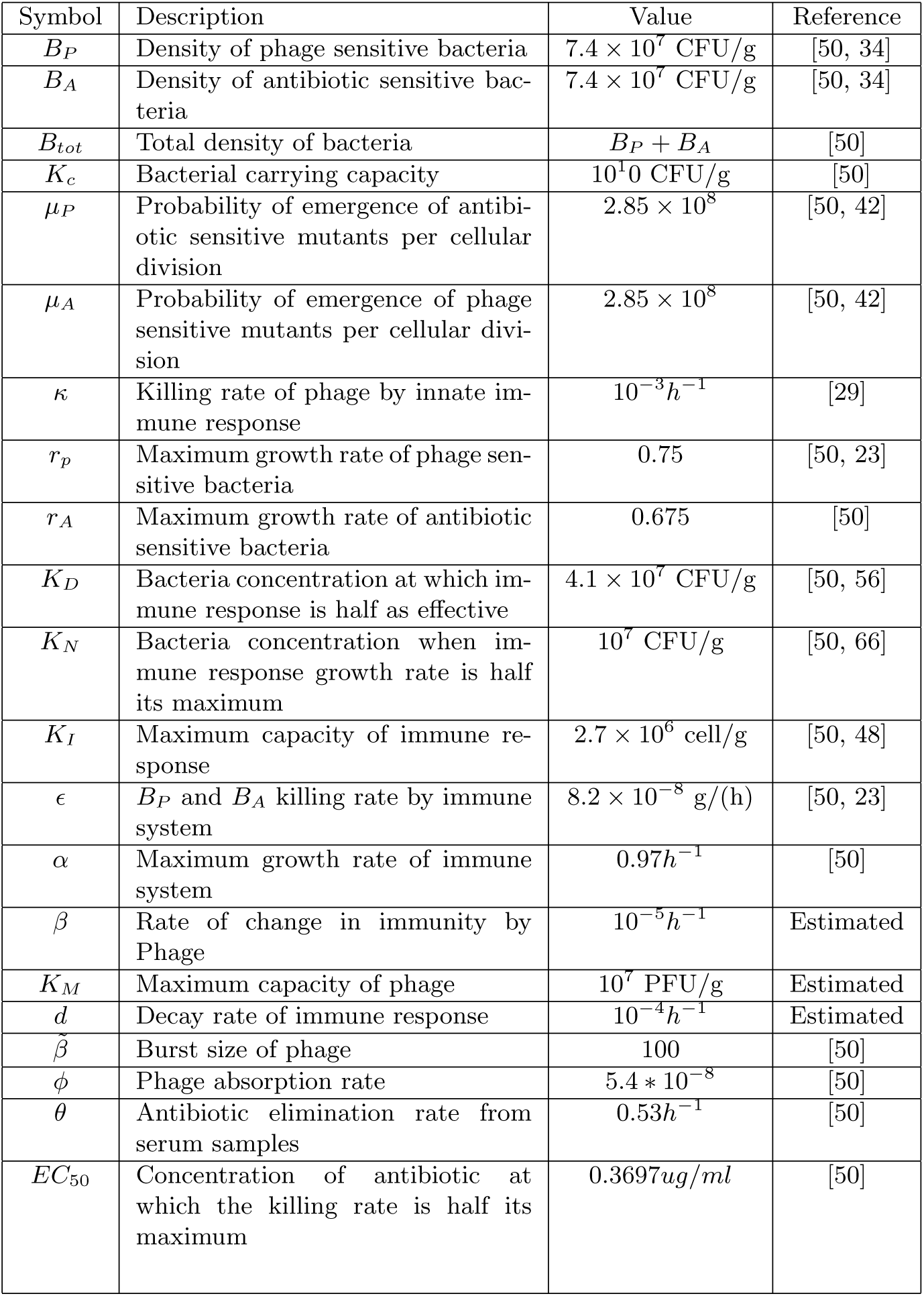
Parameters and Descriptions

| Symbol | Description | Value | Reference |
| --- | --- | --- | --- |
| $B_P$ | Density of phage sensitive bacteria | $7.4 \times 10^7$ CFU/g | [50, 34] |
| $B_A$ | Density of antibiotic sensitive bacteria | $7.4 \times 10^7$ CFU/g | [50, 34] |
| $B_{tot}$ | Total density of bacteria | $B_P + B_A$ | [50] |
| $K_c$ | Bacterial carrying capacity | $10^{10}$ CFU/g | [50] |
| $\mu_P$ | Probability of emergence of antibiotic sensitive mutants per cellular division | $2.85 \times 10^8$ | [50, 42] |
| $\mu_A$ | Probability of emergence of phage sensitive mutants per cellular division | $2.85 \times 10^8$ | [50, 42] |
| $\kappa$ | Killing rate of phage by innate immune response | $10^{-3}h^{-1}$ | [29] |
| $r_p$ | Maximum growth rate of phage sensitive bacteria | 0.75 | [50, 23] |
| $r_A$ | Maximum growth rate of antibiotic sensitive bacteria | 0.675 | [50] |
| $K_D$ | Bacteria concentration at which immune response is half as effective | $4.1 \times 10^7$ CFU/g | [50, 56] |
| $K_N$ | Bacteria concentration when immune response growth rate is half its maximum | $10^7$ CFU/g | [50, 66] |
| $K_I$ | Maximum capacity of immune response | $2.7 \times 10^6$ cell/g | [50, 48] |
| $\epsilon$ | $B_P$ and $B_A$ killing rate by immune system | $8.2 \times 10^{-8}$ g/(h) | [50, 23] |
| $\alpha$ | Maximum growth rate of immune system | $0.97h^{-1}$ | [50] |
| $\beta$ | Rate of change in immunity by Phage | $10^{-5}h^{-1}$ | Estimated |
| $K_M$ | Maximum capacity of phage | $10^7$ PFU/g | Estimated |
| $d$ | Decay rate of immune response | $10^{-4}h^{-1}$ | Estimated |
| $\tilde{\beta}$ | Burst size of phage | 100 | [50] |
| $\phi$ | Phage absorption rate | $5.4 \times 10^{-8}$ | [50] |
| $\theta$ | Antibiotic elimination rate from serum samples | $0.53h^{-1}$ | [50] |
| $EC_{50}$ | Concentration of antibiotic at which the killing rate is half its maximum | $0.3697ug/ml$ | [50] |

## 3. Analytical results

Here, we analytically explore possible treatment outcomes via equilibria analysis.

In Table 2, we summarize possible equilibria of the system, suggesting that infection dynamics can be resulted in any of the following cases under combination treatment:

I. Combination treatment fails.
II. Partial success is gained, since antibiotic sensitive bacteria die out as a result of combination (or only drug treatment). Yet, the equilibria analysis, detailed below, suggests that there might be up to three outcomes, indicating that the system might have bistable dynamics; i.e., the treatment outcomes might depend on the initial bacteria density, treatment doses and timing (see numerical results section).
III. Successful treatment. Both phage and antibiotic-sensitive bacteria get cleared.
IV. Phage treatment completely fails. It decays before clearing the phage-sensitive bacteria.
V. Phage treatment fails, yet drug treatment successfully eradicates the antibiotic sensitive bacteria.
VI. Drug treatment fails, yet phage therapy eradicates phage sensitive bacteria.

**Table 2:**
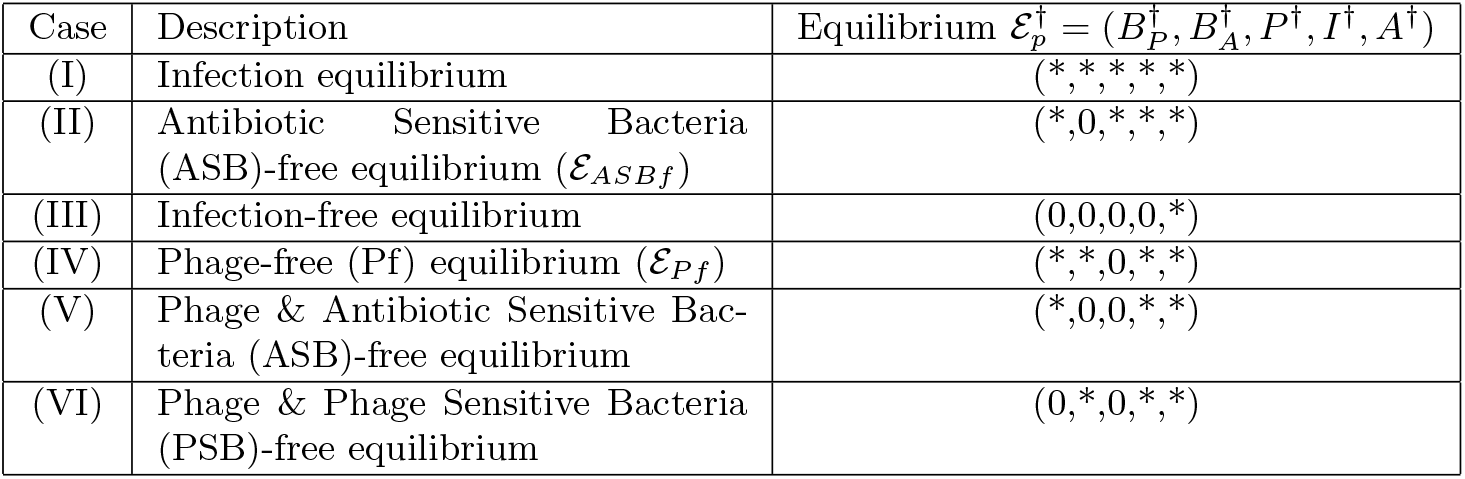
Possible equilibria of the system

A rigorous mathematical analysis and feasibility of these outcomes require stability analysis. We provide the detailed analysis of some of the cases. Below, we provide the details from the analysis of Case (II). Cases (IV) and (V) are detailed in Appendix. Due to complexity of the system, we explore the possible outcomes derived here using numerical experiments in Section 5.

### Case II. Antibiotic Sensitive Bacteria (ASB)-free equilibrium (*ε*_*ASBf*_)

In the absence of antibiotic-sensitive (phage-resistant) bacteria, we obtain the following subsystem:

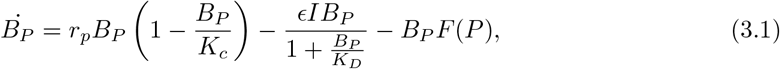

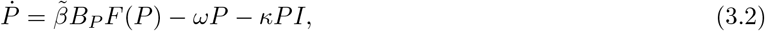

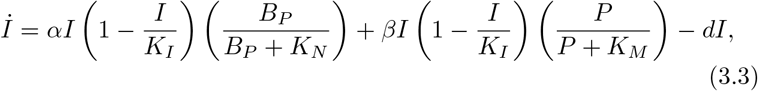

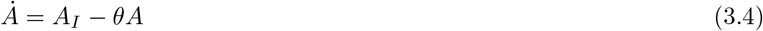

where *F* (*P*) = *ϕP* and *B*_*A*_ = 0. Equilibria of the system are the time-independent solutions. Here we are interested in phage treatment only, i.e., coexistence equilibrium

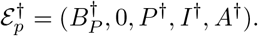

By setting the left hand of the system equal to zero, from the first equation, we obtain,

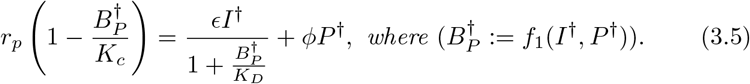

By the second equation, we also have

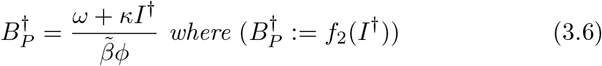

In addition, by the third equation, we get

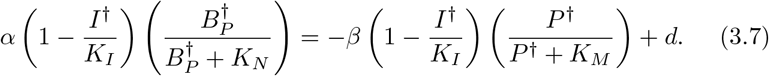

Rearranging the equality (3.7), we have

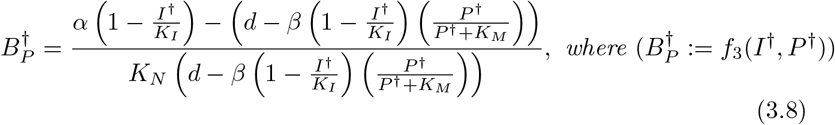

By the equality (3.8), we also have

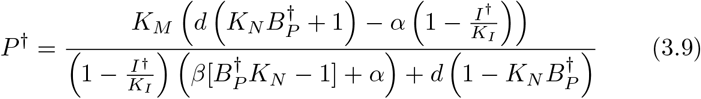

Substituting (3.9) into (3.5), we get

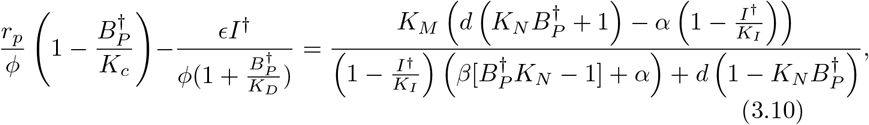

where 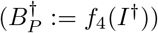.

Finally, by substituting the the right hand side of the equation (3.6) into (3.10), we get the following equality as a function of immune equilibrium component, *I*^†^:

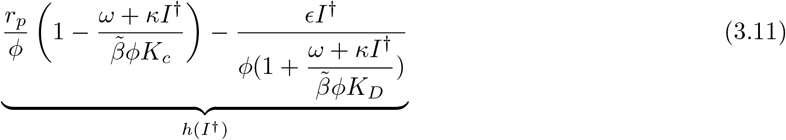

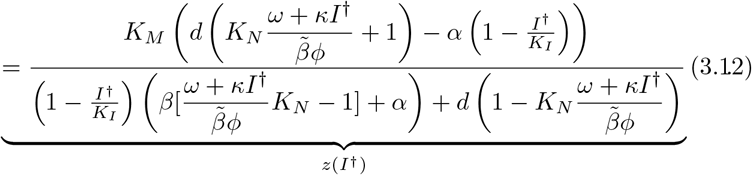

The positive intersections of the functions *h*(*I*^†^), and *z*(*I*^†^) provide the possible immune equilibrium component, *I*^†^. Notice that the left hand side of the equality, *h*(*I*^†^), is a decreasing function of *I*^†^. Moreover, the function *z*(*I*^†^) has a unique zero:

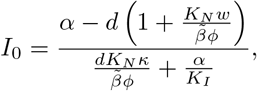

and two asymptotes *I*_1,2_:

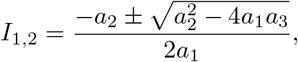

where

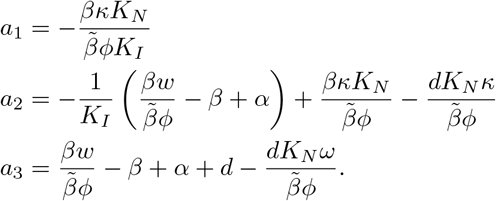

Under distinct cases with respect to sign and order of the critical points *I*_0,1,2_, the subsystem (3.1) might have zero or up to three possible positive equilibria. Note that whenever *I*^†^ > 0, we have 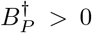 by the equation (3.6). Therefore we are looking for immune equilibrium component, *I*^†^: *I*^†^ > 0 ⇒ *P* ^†^ > 0. This result indicates that the system might have bistable dynamics; i.e., the treatment outcomes might depend on the initial bacteria/phage density, treatment doses and timing (see Section 5).

## 4. Sensitivity Analysis

Building on the work in [50], we adopt many parameter values from literature estimates and behavior fitting. However, several of our parameter values are not experimentally measurable. To determine the relative effect of fluctuations in parameter values on the model output, we use Matlab and Simbiology to implement the model and run a sensitivity analysis (similar to the process described in [55]). Sensitivity analysis of parameters for our model will inform us about changes to which parameters would have the most affect on the model transients. The following general steps were performed to produce a global sensitivity analysis for all the parameters over the simulation time period.

First, we established a set of reasonable parameters. The model needs to start at an admissible point in parameter space. We then used this fitted model to generate the discretized sensitivity matrix *S*. We then used *S* to rank parameters by sensitivity and set a threshold such that parameters with sensitivity below the threshold (insensitive) are fixed and parameters with sensitivity above the threshold (sensitive) are explored.

To apply this process to our model, we used the referenced values as a starting point as listed in Table 2. Most of these parameter values were used in [50] and we estimated the new parameter values for the full model to achieved biologically reasonable transient output for the model. All four observable model outputs (*B*_*A*_, *B*_*P*_, *P*, and *I*) were sampled at 10 time points (16 hours, 20 hours, 24 hours, 40 hours, 48 hours, and days 3-7). Given that there are 18 model parameters explored, a 40 × 18 discretized sensitivity matrix *S* is produced.

Next, we ranked the impact of each parameter on all four observable model outputs (*B*_*A*_, *B*_*P*_, *P*, and *I*) by calculating a root mean square sensitivity measure, as defined in Brun et al. [12]. For each column *j* of the normalized sensitivity matrix, we get

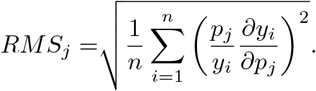

Parameter *j* is deemed insensitive if *RMS*_*j*_ is less than 5% of the value of the maximum *RMS* value calculated over all parameters. By this measure, 12 parameters were deemed insensitive, as shown in Figure 3, and fixed at their nominal values in later investigations.

**Fig. 3:**
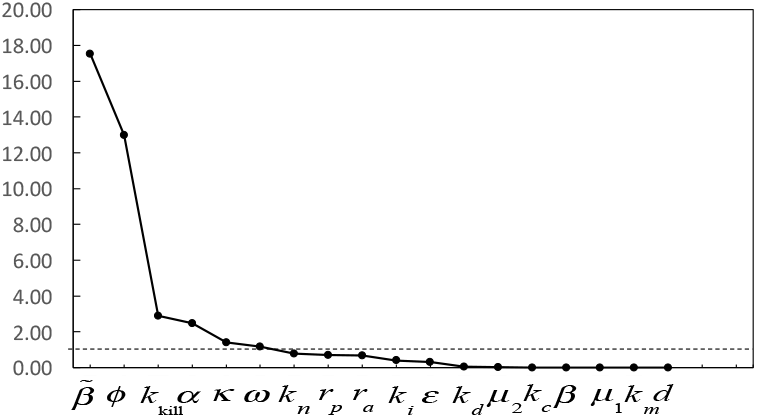
Relative sensitivities. Values below 5% of the maximum sensitivity value (indicated with the dashed line) are considered insensitive.

The strongly sensitive parameters are 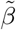 and *ϕ*. These modulate the rate of phage replication in the phage equation and also the burst rate of phage infected bacteria. Since 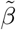 only appears in the *P* equation and it actually multiplies *ϕ*, we explored the effect of *ϕ* on the model outcome. Although not deemed as sensitive, we also chose to explore the effect of *κ*, the rate of removal of phage by immune cells, on the model outcomes since it is a new parameter in our extended system.

In the numerical results below, we explore the changes to model transients that result from different choices of these sensitive parameters.

## 5. Numerical Results

In this section, we explore numerically computed transients for some biological relevant cases of the system and apply our proposed model to investigate the interactions between bacteria, phages, antibiotics and the immune system.

### Exploring the immune response

Without any treatments (either phage or antibiotic), Fig. 4 shows that the activated immune response can clear bacteria when bacterial densities (cell densities) are low enough; however, when bacterial densities are sufficiently high, the immune system cannot mount a sufficient response to clear the infection. The complicated role of the immune response in therapeutic application of phage and antibiotics are still overgeneralize here and will be further expanded upon in later version of the model.

**Fig. 4:**
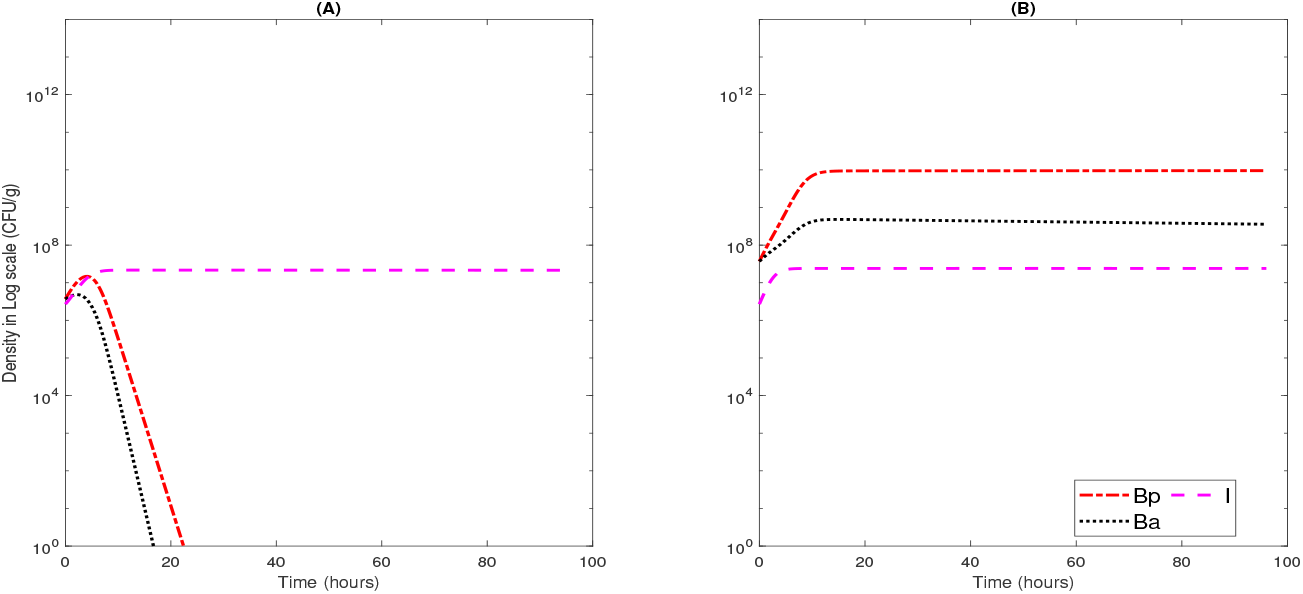
Model simulations with two levels of initial cell density. (A) a low initial cell density *B*_*P*_ (0) = *B*_*A*_(0) = 1/2 * 7.4 * 10^7^ CFU/g; (B) a high initial cell density *B*_*P*_(0) = *B*_*A*_(0) = 1/2 * 7.4 * 10^8^ CFU/g. No treatments are applied and other parameter values are fixed in Table 1.

In this model, we have included terms to allow immune cells act against the phages during the treatment [29]. This is a relevant inclusion to the model because it helps explain instances of phage ineffectiveness. One component of the new model terms, is the parameter *κ*, which describes the clearance of phages by the host immune system. Because this is a new addition to the model, we investigate the effect the value of *κ* has on the effectiveness of the combination phage-antibiotic therapy.

In Fig. 5, we use three different values of *κ*, with other parameter values fixed in Table 1 and initial bacterial levels *B*_*P*_ (0) = *B*_*A*_(0) = 1/2 * 7.4 * 10^8^ CFU/g. Also during our experiments, both phage and antibiotic therapy are received after 2 hours of the infection (*P* = 7.4 * 10^8^ PFU/g, *A* = 0.035 *ug/ml*). As can be seen in Fig. 5(A)–(C), our results show that higher values of *κ*, the killing rate of phages by immune response, results in lower availability of phages at the equilibrium state, but that the final patient outcome is not different.

**Fig. 5:**
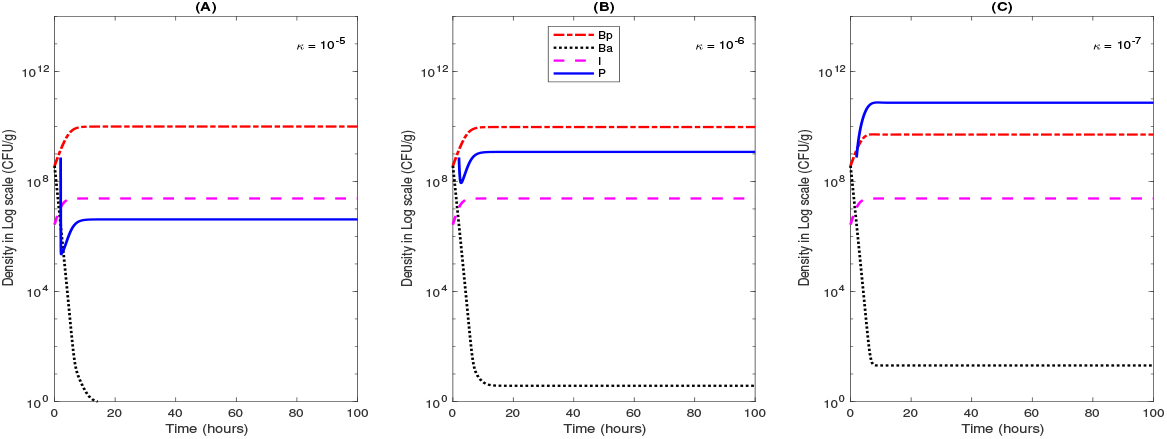
Model behaviors with different levels of killing rate, *κ*, of phage by immune cells. (A) *κ* = 10^−5^; (B) *κ* = 10^−6^; (C) *κ* = 10^−7^. Here, *B*_*P*_ (0) = *B*_*A*_(0) = 1/2 * 7.4 * 10^8^ CFU/g, both phage and antibiotic treatment were administered after 2 hours of the infection, and other parameter values fixed in Table 1.

### Effect of nonlinear phage absorption rate *ϕ*

In our simulations, we assume that phage infects and lyses *B*_*P*_ bacteria at a rate *F*(*P*), where the function *F*(*P*) = *ϕP*^0.6^ is used to account for heterogeneous mixing. In the above sensitivity analysis shown in Fig. 3, the system’s transients are sensitive to the nonlinear phage absorption rate *ϕ*. We therefore have explored the predicted effectiveness of phage therapy, as it changes with altering *ϕ*. In Fig. 6, we have shown transients for three different choices of *ϕ*, with other parameter values fixed in Table 1 and initial bacterial levels *B*_*P*_(0) = *B*_*A*_(0) = 1/2 * 7.4 * 10^8^ CFU/g. In all panels, both phage and antibiotic therapy are received 2 hours after the start of the simulation. In Fig. 6(A), where phage absorption rate is 2*ϕ*, the *B*_*A*_ goes to near zero but *B*_*P*_ stays high; while in Fig. 6(B), where phage absorption rate is 2.5*ϕ*, the same initial dose of phages is able to bring down the level of *BP* to zero, and we attain the trivial equilibrium; while in Fig. 6(C), where phage absorption rate is 5*ϕ*, the *B*_*P*_ goes to zero and the process occurs faster than compared to (B).

**Fig. 6:**
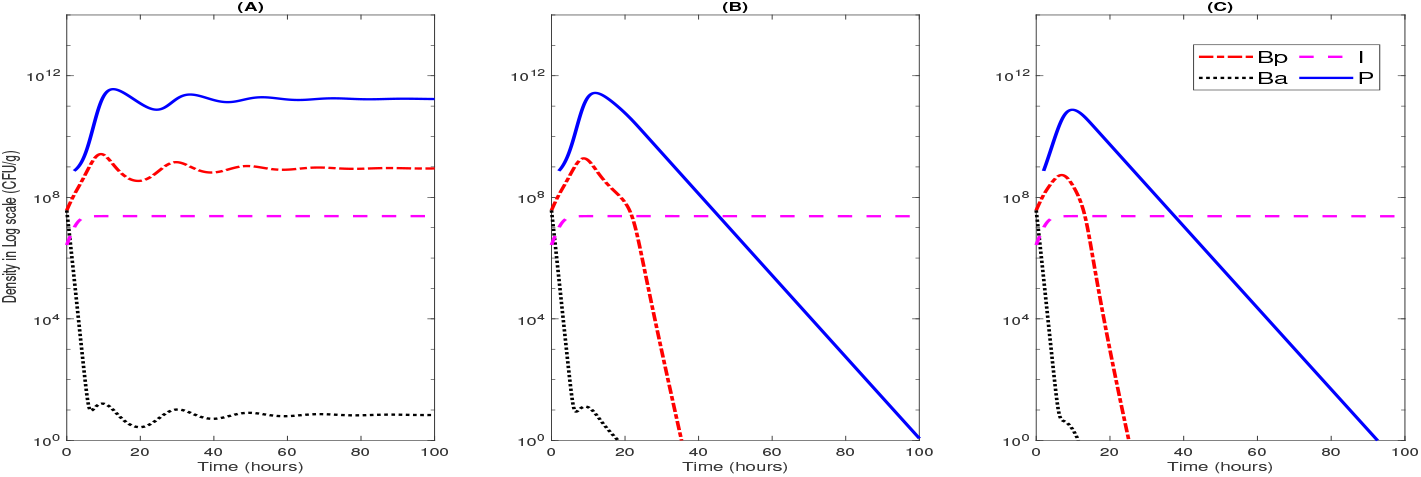
Model behaviors with different levels of phage absorption rate. (A) *ϕ* = 2 * 5.4 * 10^−8^; (B) *ϕ* = 2.5 * 5.4 * 10^−8^; (C) *ϕ* = 5 * 5.4 10^−8^. Here, *B*_*P*_(0) = *B*_*A*_(0) = 1/2 * 7.4 * 10^8^ CFU/g, both phages and antibiotic therapy are administered after 2 hours of the infection, and other parameter values fixed in Table 1.

### Effect of time of administration of phage dose

Next, we investigate the effects of timing of phage therapy on the outcome of the infection. In all the simulations, antibiotics are given at the start of the simulation, the initial bacterial levels are *B*_*P*_(0) = *B*_*A*_(0) = 1/20 * 7.4 * 10^8^ CFU/g (a relatively low level), the nonlinear phage absorption rate is 2*ϕ*, and the other parameter values are fixed as shown in Table 1. In Fig. 7(A), the phage dose is given 2 hours after infections. We can see that the antibiotic-sensitive bacteria, *B*_*A*_, decays quickly and goes to equilibrium near zero. Even though phage therapy lowers the *B*_*P*_ bacteria, *B*_*P*_ does not get completely removed from the system and a non-zero equilibrium is achieved for both *B*_*A*_ and *B*_*P*_. However, in Fig. 7(B), the phage is administered after 10 hrs after the start of infection: the density of *B*_*P*_ is already high, which helps the phages to grow and in turn phages are able to reduce the density of *B*_*P*_. Then in the absence of *B*_*P*_, the phage level also goes to zero. These experiments indicates that the timing of phage therapy can be an important factor because phage effectiveness depends on the density of bacteria present in the system.

**Fig. 7:**
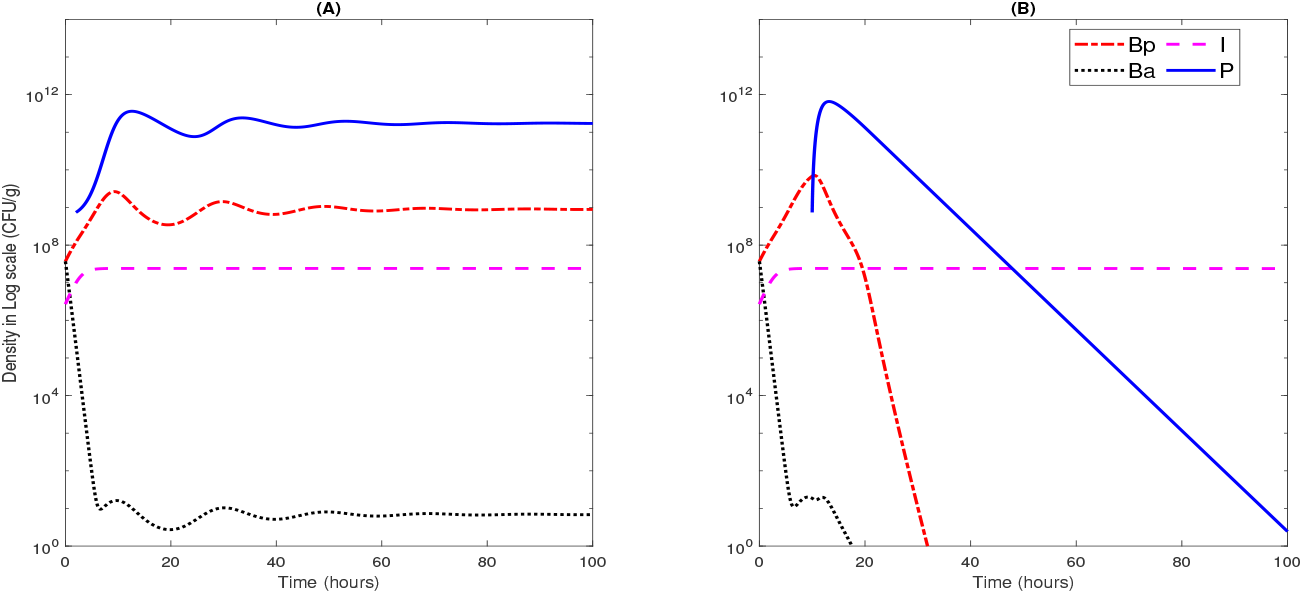
Model behaviors with different timing of phage therapy. (A) Phage dose 7.4 * 10^8^ PFU/g was administered after 2 hours; (B) phage dose 7.4 * 10^8^ PFU/g was administered after 10 hrs from the beginning of infection. Here, the initial bacterial level is *B*_*P*_ (0) = *B*_*A*_(0) = 1/20 7.4 * 10^8^ CFU/g (a relatively low level), the nonlinear phage absorption rate is 2*ϕ*, and the other parameter values are fixed as shown in Table 1.

### Varying time and quantity of phage dose in multi-dose regi-men

We continue our experiments by varying the frequency and quantity of the phage therapy dose to explore possible outcomes of phage therapy. For both simulations in Fig. 8 the same initial infection level *B*_*P*_(0) = *B*_*A*_(0) = 1/2 * 7.4 * 10^8^ CFU/g are used, the same antibiotic therapy is administered after 2 hours, and the parameter values are same as in Table 1. In the first experiment (Fig. 8(A)), we use only one dose of phage. The dose of phage (*P* = 7.4 * 10^8^ PFU/g) is administered after 2 hours of infection. It is shown that the *B*_*A*_ goes to nearly zero, but *B*_*P*_ goes to a positive equilibrium (*B*_*P*_ ≫ 0). That is, we do not have a successful treatment. In the second experiment, we explore two doses of phage. As in (Fig. 8(A)), the first dose (*P* = 7.4 * 10^8^ PFU/g) is administered after 2 hours of infection. Then 10 hours after the first dose, the second dose (*P* = 2.4 * 10^12^ PFU/g) is given. We found that if the amount of second dose of phage is high enough, then the density of *B*_*P*_ goes to zero rapidly, and we obtain a successful treatment at the end. Otherwise, you need to do more doses of phage treatment (See Fig. 9).

**Fig. 8:**
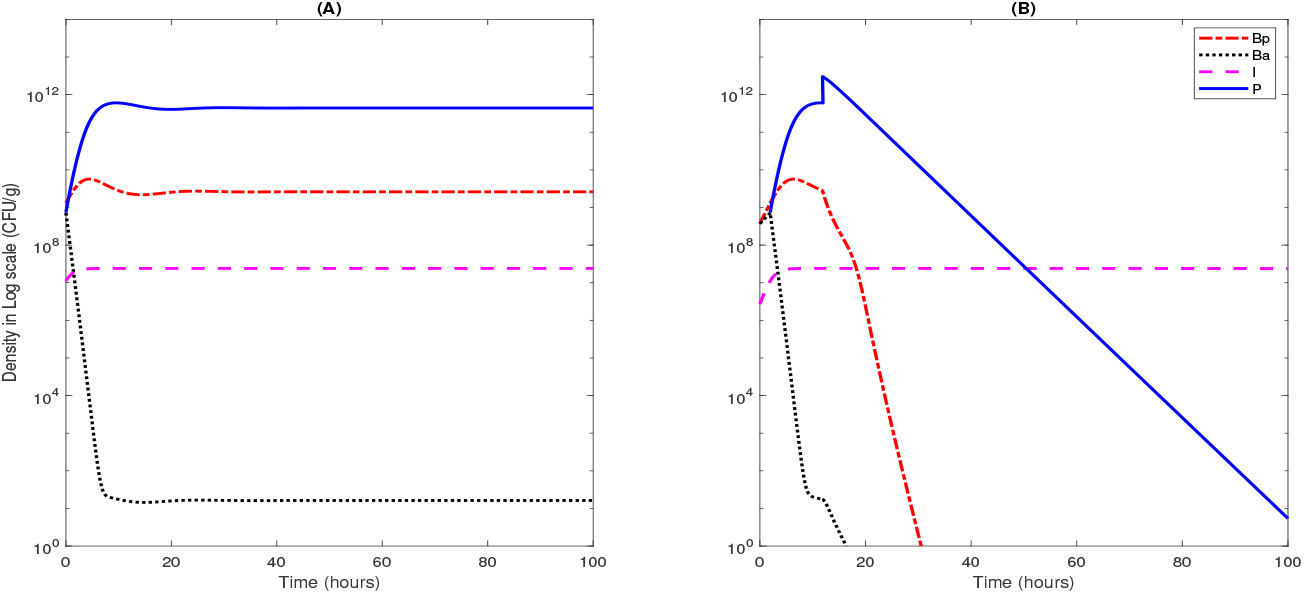
Model behavior in multi-dose regimen of phage therapy. (A) One dose of phage treatment. The only dose 7.4 * 10^8^ PFU/g was administered after 2 hours of the beginning of infection; (B) two doses of phage treatment. The first dose 7.4 * 10^8^ PFU/g was administered after 2 hours of the beginning of infection and the second dose *P* = 2.4 10^12^ PFU/g was administered after 10 hours of first dose. Here, the initial bacterial level *B*_*P*_(0) = *B*_*A*_(0) = 1/2 * 7.4 * 10^8^ CFU/g is used, antibiotic therapy is administered after 2 hours, and parameter values are fixed as shown in Table 1.

**Fig. 9:**
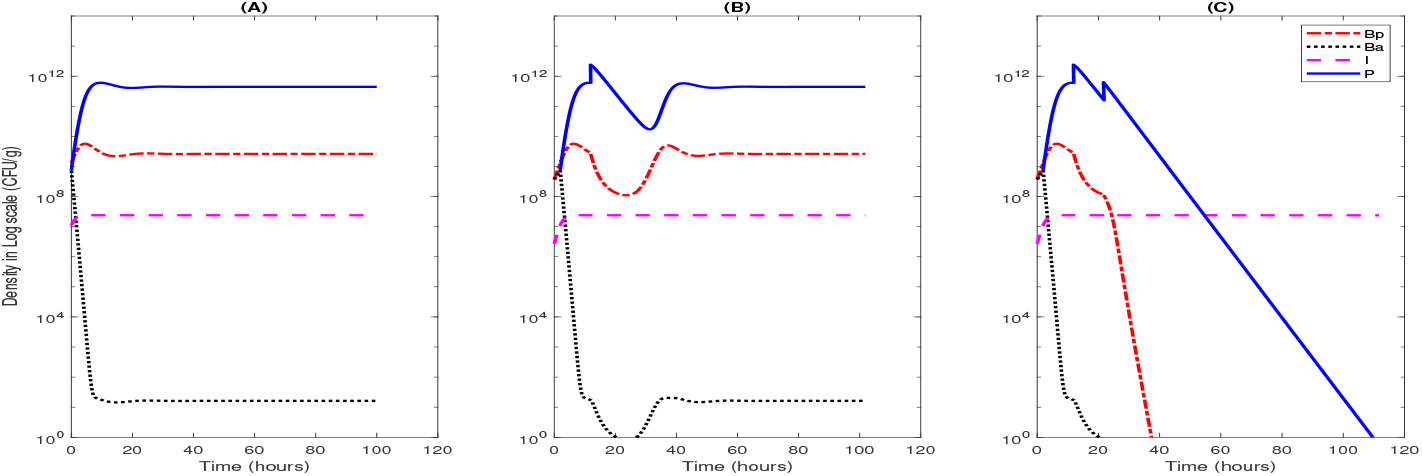
Model behavior in multi-dose regimen of phage therapy. (A) One dose of phage treatment. The only dose 7.4 * 10^8^ PFU/g was administered after 2 hours of the beginning of infection; (B) two doses of phage treatment. The first dose 7.4 * 10^8^ PFU/g was administered after 2 hours of the beginning of infection and the second dose *P* = 1.8 * 10^12^ PFU/g was administered after 10hrs of first dose; (C) three doses of phage treatment. The first dose 7.4 * 10^8^ PFU/g was administered after 2 hours of the beginning of infection, the second dose *P* = 1.8 * 10^12^ PFU/g was administered after 10hrs of first dose, and the third dose *P* = 4.5 * 10^11^ PFU/g was administered after 10hrs of second dose. Here, the initial bacterial level *B*_*P*_(0) = *B*_*A*_(0) = 1/2 * 7.4 * * 10^8^ CFU/g is used, antibiotic therapy is administered after 2 hours, and parameter values are fixed as shown in Table 1.

In Fig. 9, three experiments are shown. In all simulations, the same initial infection level *B*_*P*_(0) = *B*_*A*_(0) = 1/2 * 7.4 * 10^8^ CFU/g is used, antibiotic therapy is administered after 2 hours, and parameter values are fixed as shown in Table 1. To conduct the comparison study, the first experiment (Fig. 9(A)), is same as Fig. 8, i.e., only one dose of phages (*P* = 7.4 * 10^8^ PFU/g) is administered after 2 hours of infection and it did not lead to a successful treatment. In the second experiment (Fig. 9(B)), we administer two doses of phages. The first dose (*P* = 7.4 * 10^8^ PFU/g) is administered after 2 hours of infection. Then 10 hours after the first dose, the second dose (*P* = 1.8 * 10^12^ PFU/g, a relatively low value compared to 8(B)) is given. We found that even though the *B*_*P*_ density decreases quickly after the second dose of phage, it eventually rebound. This indicates we still fail the treatment. In the third experiment (Fig. 9(C)), we have three doses of phage therapy. The first two doses are administered as in the second experiment, i.e., the first dose (*P* = 7.4 * 10^8^ PFU/g) is administered after 2 hours of infection. Then 10 hours after the first dose, the second dose (*P* = 1.8 10^12^ PFU/g, a relatively low value compared to 8(B)) is given. Now, 10 hours after the second dose, we try the third dose (*P* = 4.5 * 10^11^ < 1.8 * 10^12^ PFU/g), and see that we can obtain a successful treatment. Hence, we believe that the number of doses and the size of the dose of phages have significant impacts on the clearance of the bacterial infection.

## 6. Discussion

In this work, we have analyzed a prior model, developed in [50], for the use of combination antibiotic and phage therapy for the treatment of a systemic bacterial infection. We extended it by including immune response to circulating phages. While phages are not “infectious” to humans, they are a foreign substance in the body and will elicit an inflammatory response. Additionally, the innate immune response of the patient will clear some of the phages either through filtration or through phagocytosis. Therefore, we would like to see if this dynamic is important to consider for predicting the effectiveness of the combination therapy.

By utilizing equilibria analysis, we find that the model proposed by Rodriguez-Gonzalez, et al. [50] can have six possible steady-state cases for model outcomes. In addition, our analysis suggests that in some cases, the system might display bistable dynamics; i.e., the treatment outcomes can depend on initial conditions, determined by dose of drug or phage cocktail, or timing of any of these treatments, or the frequency of these treatments in combinations. Therefore, we numerically explores outcomes of treatment options using phage therapy in combination with antibiotic treatment in order to gain insights of how to optimize the treatment outcomes.

We performed a sensitivity analysis to determine which parameters are likely to affect the transient behavior and the overall outcome of the system. To that end, we found that two parameters were of the most interest. The one with the most biological meaning (*ϕ*) was investigated and found to have an effect on the outcome of the system. It will be important in future modeling work to better estimate the number of phage released during lysis while in a human host.

The timing of the phage treatment was also important for determining patient outcome. Because phages replicate inside the bacteria, the level of bacterial infection at the time of treatment initiation influences the effectiveness of the phage therapy. Repeated dosing with phages is also helpful in clearing the bacterial infection. Determining dosing protocols and quantifying the related risks will be important for future studies.

This initial investigation has been fruitful for understanding some of the competing dynamics observed in antibiotic/phage combination therapy, and has opened the work up to further questions and lines of research:

- Are there further interactions with the host immune system that need to be explored? (Innate/adaptive/filtering)
- Can we determine an optimal treatment strategy?
- Do different bacterial infections require different parameter values or are there other considerations that need to be made? Some bacteria have “broad spectrum” response to phages and some require treatment with very specific phages.
- How fast do bacteria develop or lose immunity to phages?
- What additional complications occur in immunocompromized individuals?

There is hope that phage therapy will usher in a new line of treatment for difficult bacterial infections but there are much work needed to understand the complex dynamics and to devise effective, broadly implementable treatment protocols.

## Acknowledgements

The work described herein was initiated during the Collaborative Workshop for Women in Mathematical Biology hosted by the Institute for Pure and Applied Mathematics at the University of California, Los Angeles in June 2019. Funding for the workshop was provided by IPAM, the Association for Women in Mathematics’ NSF ADVANCE “Career Advancement for Women Through Research-Focused Networks” (NSF-HRD 1500481) and the Society for Industrial and Applied Mathematics. The authors thank the organizers of the IPAM-WBIO workshop (Rebecca Segal, Blerta Shtylla, and Suzanne Sindi) for facilitating this research.

## APPENDIX

## Case IV. Phage-free equilibrium 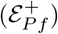

Setting *P* = 0, at the steady-state we obtain the following equation system:

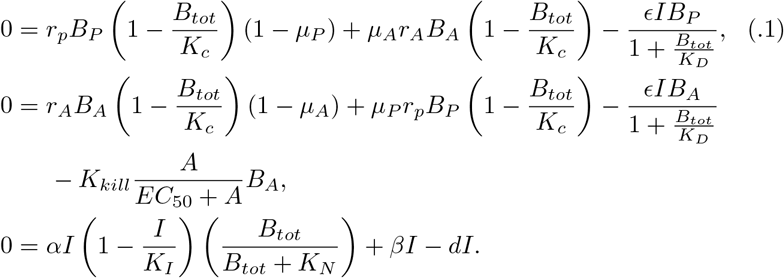

By the last equation in (.1), we have

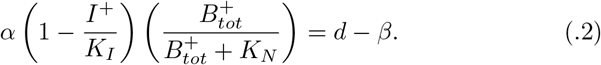

Then rearranging (.2), we obtain

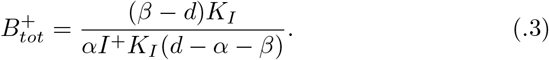

Also by the first and second equations in (.1), we obtain

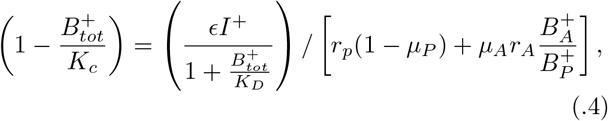

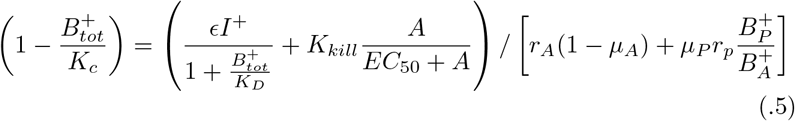

By the equality of equations in (.4), we have

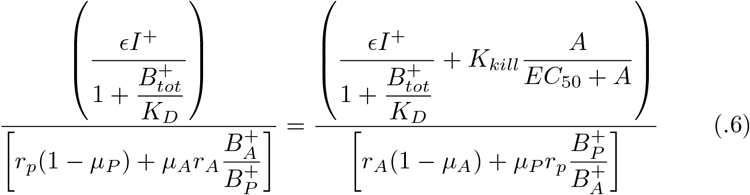

Let 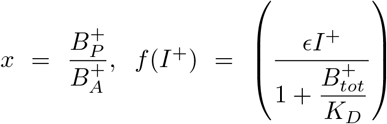, *a*_0_ = *r*_*p*_(1 − *μ_P_*), 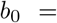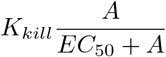, *c*_0_ = *r*_*A*_(1 − *μ*_*A*_), *a*_1_ = *μ*_*A*_*r*_*A*_, and *c*_1_ = *μ*_*P*_*r*_*p*_. Then by (.6), we obtain

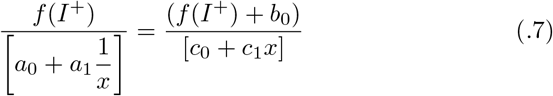

Then rearranging it, we have

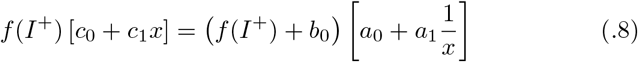

Multiplying both sides with *x* and rearranging we obtain

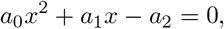

where

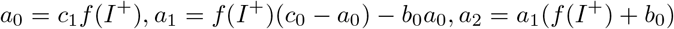

Therefore we get the steady-state ratio 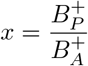 as follows:

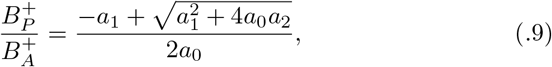

where

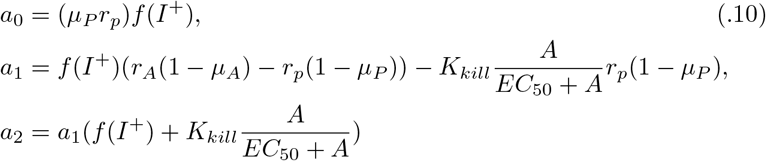

with 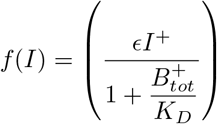 and 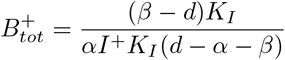.

Also note that 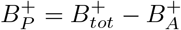. Then by (.9), we obtain

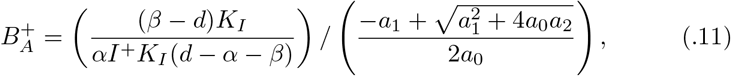

where the expressions of *a*_*i*_ for *i* = 0, 1, 2 are given in (.10). Therefore the system has at most one positive phage-free equilibrium 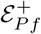.

## Case V. Phage & Antibiotic Sensitive Bacteria (ASB)-free equilibrium 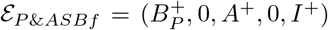

Setting *P* = *B*_*A*_ = 0, we obtain the following equation system:

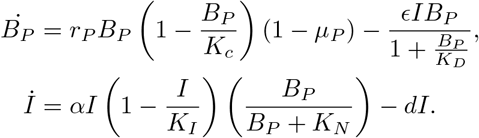

At the steady state, by the second equation, we have

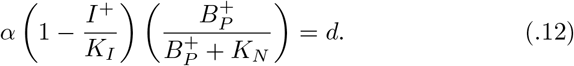

Rearranging it, we obtain

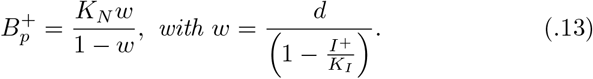

By the first equation, we also have

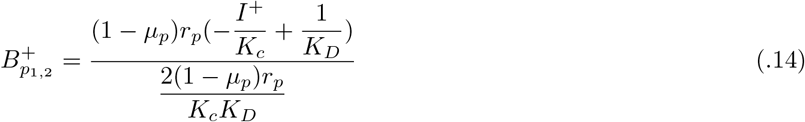

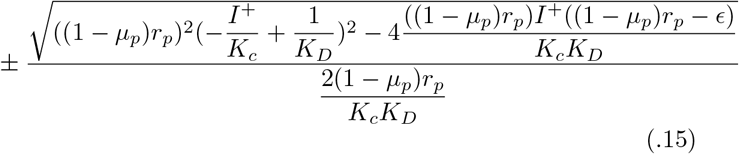

Note that the equalities (.13) and (.14) are functions of *I*^+^, and intersection of both equations give the equilibrium *I*^+^ component of the equilibria of the system, and the other component of the equilibria can be found by substituting the component *I*^+^ into the equation (.13). It is clear that the system can have more than one phage & antibiotic sensitive bacteria-free equilibrium 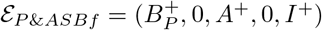.

## REFERENCES

[1] Abedon, S.: Kinetics of phage-mediated biocontrol of bacteria. Foodborne Pathog. Dis. 6(7), 807–815 (2009)

[2] Ankomah, P., Levin, B.R.: Exploring the collaboration between antibiotics and the immune response in the treatment of acute, self-limiting infections. Proceedings of the National Academy of Science. 111(23), 8331–8338 (2014)

[3] Antibiotic/antimicrobial resistance: Center for disease control and prevention (September 10, 2018)

[4] Austin, D., Anderson, R.: Studies of antibiotic resistance within the patient, hospitals and the community using simple mathematical models. Philos Trans R Soc Lond B Biol Sc. 354(1384), 721–738 (1999)

[5] Austin, D.J., Kakehashi, M., Anderson, R.M.: The transmission dynamics of antibiotic-resistant bacteria: the relationship between resistance in commensal organisms and antibiotic consumption. Philos. Trans. R. Soc. Lond. B: Biol. Sci. 264, 1629–1638 (1997)

[6] Bergstrom, C.T., Lipsitch, M., , Levin, B.R.: Natural selection, infectious transfer and the existence conditions for bacterial plasmids. Genetic. 155(4), 1505–1519 (2000)

[7] Bergstrom, C.T., Lo, M., Lipsitch, M.: The transmission dynamics of antibiotic-resistant bacteria: the relationship between resistance in commensal organisms and antibiotic consumption. Proc. Natl. Acad. Sci. USA. 101, 13,285–13,290 (2004)

[8] Bonhoeffer, S., Liptsitch, M., Levin, B.R.: Evaluating treatment protocols to prevent antibiotic resistance. Proc. Natl. Acad. Sci. US. 101, 12,106–12,111 (1996)

[9] Bootsma, M.C., Diekmann, J.O., Bonten, M.J.M.: Controlling methicillin-resistant Staphylococcus aureus: quantifying the effects of interventions and rapid diagnostic testing. Proc. Natl. Acad. Sci. US. 103, 5620–5625 (2006)

[10] Browne, C., Wang, M., Webb, G.F.: A stochastic model of nosocomial epidemics in hospital intensive care units. Electron. J. Qual. Theory Differ. Equ. 6, 1–12 (2017)

[11] Browne, C., Webb, G.F.: A nosocomial epidemic model with infection of patients due to contaminated rooms. Math Biosci Eng. 12(4), 761–787 (2015)

[12] Brun, R., Reichert, P., Künsch, H.R.: Practical identifiability analysis of large environmental simulation models. Water Resources Researc. 37(4), 1015–1030 (2001)

[13] Campbell, A.: Conditions for the existence of bacteriophage. Evolutio. 15(2), 153–165 (1961)

[14] Chamchod, F., Ruan, S.: Modeling methicillin-resistant Staphylococcus aureus in hospitals: transmission dynamics, antibiotic usage and its history. Theor. Biol. Med. Model. 9, 25 (2012)

[15] Chan B.k. and Sistrom, M., Wertz, J., Kortright, K., Narayan, D., Turner, P.: Phage selection restores antibiotic sensitivity in mdr Pseudomonas aeruginosa. Scientific Reports 6(26717) (2016)

[16] Chao, L., Levin, B., Stewart, F.: A complex community in a simple habitat: an experimental study with bacteria and phage. Ecolog. 58(2), 369–378 (1977)

[17] Clark, J.: Bacteriophage therapy: history and future prospects. Future Virol. 10(4), 449–461 (2015)

[18] Cooper, B., Medley, G., Stone, S., Kibbler, C., Cookson, B., Roberts, J., Duckworth, G., Lai, R., Ebrahim, S.: methicillin-resistant Staphylococcus aureus in hospitals and the community: stealth dynamics and control catastrophes. Proceedings of the National Academy of Sciences of the United States of Americ. 101(27), 10,223–10,228 (2004)

[19] D’Agata, E.M., Horn, M.A., Ruan, S., Webb, G.F., Wares, J.R.: Efficacy of infection control interventions in reducing the spread of multidrug-resistant organisms in the hospital setting. PLoS On. 7(2), e30,170 (2012)

[20] D’Agata, E.M.C., Webb, G.F., Horn, M.A.: A mathematical model quantifying the impact of antibiotic exposure and other interventions on the endemic prevalence of vancomycin-resistant Enterococci. J. Infect. Disease. 192, 2004–2011 (2005)

[21] Davies, O.L.: Who publishes list of bacteria for which new antibiotics are urgently needed (2017). URL https://www.who.int/news-room/detail/27-02-2017-who-publishes-list-of-bacteria-for-which-new-antibiotics\-are-urgently-needed

[22] Dennehy, J.: What can phages tell us about host-pathogen coevolution? Int. J. Evol. Biol. 2012(396165), 12 (2012)

[23] Drusano, G.L., Vanscoy, B., Liu, W., Fikes, S., Brown, D., Louie, A.: Saturability of granulocyte kill of pseudomonas aeruginosa in a murine model of pneumonia. Antimicrobial agents and chemotherap. 55, 2693–2695 (2011)

[24] DAgata, E.M., Magal, P., Olivier, D., Ruan, S., Webb, G.F.: Modeling antibiotic resistance in hospitals: the impact of minimizing treatment duration. Journal of Theoretical Biolog. 249(3), 487–499 (2007)

[25] F. Jafri, H., Bonten, M.: Alternatives to antibiotics. Intensive Care Medicin. 42(12), 2034–2036 (2016)

[26] Guenther, S., Huwyler, D., Richard, S., Loessner, M.: Virulent bacteriophage for efficient biocontrol of listeria monocytogenes in ready-to-eat foods. Appl. Environ. Microbiol. 75(1), 93–100 (2009)

[27] Hagens, S., Loessner, M.: Application of bacteriophages for detection and control of foodborne pathogens. Appl. Microbiol. Biotechnol. 76(3), 513–519 (2007)

[28] Hall, I.M., Barrass, I., Leach, S., Pittet, D., Hugonnet, S.: Transmission dynamics of methicillin-resistant Staphylococcus aureus in a medical intensive care unit. Journal of the Royal Society, Interfac. 9(75), 2639–2652 (2012)

[29] Hodyra-Stefaniak, K., Miernikiewicz, P., Drapa-la, J., Drab, M., Jończyk-Matysiak, E., Lecion, D., Kázmierczak, Z., Beta, W., Majewska, J., Harhala, M., Bubak, B., K-lopot, A., Górski, A., Dąbrowska, K.: Mammalian host-versus-phage immune response determines phage fate in vivo. Scientific Report. 5, 14,802 (2015)

[30] Huang, Q., Horn, M.A., Ruan, S.: Modeling the effect of antibiotic exposure on the transmission of methicillin-resistant staphylococcus aureus in hospitals with environmental contamination. Mathematical Biosciences and Engineerin. 16(5), 3641–3673 (2019)

[31] Huang, Q., Huo, X., Miller, D., Ruan, S.: Modeling the seasonality of methicillin-resistant Staphylococcus aureus infections in hospitals with environmental contamination. J Biol Dyn. 13(sup1), 99–122 (2018)

[32] Huang, Q., Huo, X., Ruan, S.: Optimal control of environmental cleaning and antibiotic prescription in an epidemiological model of methicillin-resistant Staphylococcus aureus infections in hospitals. Math. Biosci. 311, 13–30 (2019)

[33] Kingwell, K.: Bacteriophage therapies re-enter clinical trials. Nature Reviews Drug Discover. 14(8), 515–516 (2015)

[34] Leung, C.Y.J., Weitz, J.S.: Modeling the synergistic elimination of bacteria by phage and the innate immune system. Journal of theoretical biolog. 429, 241–252 (2017)

[35] Levin, B., Stewart, F., Chao, L.: Resource-limited growth, competition, and predation: a model and experimental studies with bacteria and bacteriophage. Am. Nat. 111(977), 3–24 (1977)

[36] Levin, B.R., Bergstrom, C.T.: Bacteria are different: observations, interpretations, speculations, and opinions about the mechanisms of adaptive evolution in prokaryotes. Proceedings of the National Academy of Science. 97(13), 6981–6985 (2000)

[37] Levin, B.R., Bull, J.J.: Population and evolutionary dynamics of phage therapy. Nature Reviews Microbiolog. 2(2), 166–173 (2004)

[38] Levin, B.R., Stewart, F.M.: The population biology of bacterial plasmids: a priori conditions for the existence of mobilizable nonconjugative factors. Genetic. 94(2), 425–443 (1980)

[39] Lin, D.M., Koskella, B., Lin, H.C.: Phage therapy: An alternative to antibiotics in the age of multi-drug resistance. World journal of gastrointestinal pharmacology and therapeutic. 8, 162 (2017)

[40] Lipsitch, M., Bergstrom, C.T., Levin, B.R.: The epidemiology of antibiotic resistance in hospitals: paradoxes and prescriptions. Proc. Natl. Acad. Sci. US. 97, 1938–1943 (2000)

[41] Luepke, K.H., Suda, K.J., Boucher, H., Russo, R.L., Bonney, M.W., Hunt, T.D., Mohr III, J.F.: Past, present, and future of antibacterial economics: increasing bacterial resistance, limited antibiotic pipeline, and societal implications. Pharmacotherapy: The Journal of Human Pharmacology and Drug Therap. 37, 71–84 (2017)

[42] Luria, S.E., Delbrück, M.: Mutations of bacteria from virus sensitivity to virus resistance. Genetic. 28, 491 (1943)

[43] Meyer, J., Dobias, D., Weitz, J., Barrick, J., Quick, R., Lenski, R.: Repeatability and contingency in the evolution of a key innovation in phage lambda. Scienc. 335(6067), 428–432 (2012)

[44] Pincus, N., Reckhow, J., Saleem, D., Jammeh, M., Datta, S., Myles, I.: Strain specific phage treatment for staphylococcus aureus infection is influenced by host immunity and site of infection. PLoS ON. 10(4), 1–16 (2015)

[45] Potera, C.: Phage renaissance: new hope against antibiotic resistance. Environ Health Perspect. 121, a48–53 (2013)

[46] Rea, K., Dinan, T.G., Cryan, J.F.: The microbiome: A key regulator of stress and neuroinflammation. Neurobiol Stres. 4, 23–33 (2016)

[47] Reardon, S.: Phage therapy gets revitalized. Natur. 510(7503), 15–16 (2014)

[48] Reutershan, J., Basit, A., Galkina, E.V., Ley, K.: Sequential recruitment of neutrophils into lung and bronchoalveolar lavage fluid in LPS-induced acute lung injury. American Journal of Physiology-Lung Cellular and Molecular Physiolog. 289, L807–L815 (2005)

[49] Roach, D.R., Leung, C.Y., Henry, M., Morello, E., Singh, D., Di Santo, J.P., Weitz, J.S., Debarbieux, L.: Synergy between the host immune system and bacteriophage is essential for successful phage therapy against an acute respiratory pathogen. Cell host & microbe 22(1), 38–47 (2017)

[50] Rodriguez-Gonzalez, R.A., Leung, C.Y., Chan, B.K., Turner, P.E., Weitz, J.S.: Quantitative models of phage-antibiotics combination therapy. BioRxiv, 633784 (2019)

[51] Seo, J., Seo, D., Oh, H., Jeon, S., Oh, M.H., Choi, C.: Inhibiting the growth of escherichia coli O157:H7 in beef, pork, and chicken meat using a bacteriophage. Korean J. Food Sci. Anim. Resour. 2(2), 186–93 (2016)

[52] Stewart F. M., ., Levin, B.R.: The population biology of bacterial plasmids: a priori conditions for the existence of conjugationally transmitted factors. Genetic. 87(2), 209–228 (1977)

[53] Thiel, K.: Old dogma, new tricks–21st century phage therapy. Nat. Biotechnol. 22(1), 31–36 (2004)

[54] Tiwari, B., Kim, S., Rahman, M., Kim, J.: Antibacterial efficacy of lytic pseudomonas bacteriophage in normal and neutropenic mice models. J. Microbiol. 49(6), 994–999 (2011)

[55] Torres, M., Wang, J., Yannie, P., Ghosh, S., Segal, R., Reynolds, A.: Identifying important parameters in the inflammatory process with a mathematical model of immune cell influx and macrophage polarization. PLoS Comput Bio. 15(7), 497–503 (2019)

[56] Torres-Barceló, C., Franzon, B., Vasse, M., Hochberg, M.E.: Long-term effects of single and combined introductions of antibiotics and bacteriophages on populations of pseudomonas aeruginosa. Evolutionary application. 9, 583–595 (2011)

[57] Trigo, G., Martins, T., Fraga, A., Longatto-Filho, A., Castro, A., Azeredo, J., Pedrosa, J.: Phage therapy is effective against infection by mycobacterium ulcerans in a murine footpad model. PLoS Negl. Trop. Di. 7(4), e2183 (2013)

[58] Wang, J., Wang, L., Magal, P., Wang, Y., Zhuo, S., Lu, X., Ruan, S.: Modelling the transmission dynamics of Meticillin-resistant Staphylococcus aureus in Beijing Tongren Hospital. Journal of Hospital Infectio. 79(4), 302–308 (2011)

[59] Wang, L., Ruan, S.: Modeling nosocomial infections of methicillin-resistant Staphylococcus aureus with environment contamination. Scientific Reports 7(2017)

[60] Wang, X., Xiao, Y., Wang, J., Lu, X.: A mathematical model of effects of environmental contamination and presence of volunteers on hospital infections in China. Journal of Theoretical Biolog. 293, 161–173 (2012)

[61] Wang, X., Xiao, Y., Wang, J., Lu, X.: Stochastic disease dynamics of a hospital infection model. Mathematical Bioscience. 241(1), 115–124 (2013)

[62] Webb, G.F.: Individual based models and differential equations models of nosocomial epidemics in hospital intensive care units. Discrete & Continuous Dynamical Systems-Series B 22(3) (2017)

[63] Webb, G.F., D’Agata, E.M.C., Magal, P., Ruan, S.: A model of antibiotic resistant bacterial epidemics in hospitals. Proc. Natl. Acad. Sci. US. 102, 13,343–13,348 (2005)

[64] Weitz, J., Hartman, H., Levin, S.: Coevolutionary arms races between bacteria and bacteriophage. Proc. Natl. Acad. Sci. USA. 102(27), 9535–9540 (2005)

[65] World Health Organization: Antimicrobial resistance - global action plan (2015)

[66] Zhang, Z., Louboutin, J.P., Weiner, D.J., Goldberg, J.B., Wilson, J.M.: Human airway epithelial cells sense pseudomonas aeruginosa infection via recognition of flagellin by toll-like receptor 5. Infection and immunit. 73, 7151–7160 (2005)

